# Absence of atrial smooth muscle in the heart of the loggerhead sea turtle (*Caretta caretta*): a re-evaluation of its role in diving physiology

**DOI:** 10.1101/2022.08.01.502306

**Authors:** Leah M. Costello, Daniel García-Párraga, Jose Luis Crespo-Picazo, Jonathan Codd, Holly A. Shiels, William Joyce

## Abstract

Contraction of atrial smooth muscle in the hearts of semi-aquatic emydid turtles regulates ventricular filling, and it has been proposed that it could regulate stroke volume during characteristic rapid transitions in cardiac output associated with diving. For this hypothesis to be supported, atrial smooth muscle should be widely distributed in diving Testudines. To further understand the putative function and evolutionary significance of endocardial smooth muscle in Testudines, we studied the hearts of loggerhead sea turtles, *Caretta caretta* (n=7), using immunohistochemistry and histology. Surprisingly, we found no evidence of prominent atrial smooth muscle in *C. caretta*. However, smooth muscle was readily identified in the sinus venosus. Our results suggest atrial smooth muscle does not contribute to the diving capabilities of *C. caretta*, indicating that the possible roles of smooth muscle in emydid turtle hearts requires a re-evaluation. In sea turtles, the sinus venosus may instead contribute to regulate cardiac filling.

## 1. Introduction

An extensive layer of atrial smooth muscle has been demonstrated in members of the semi-aquatic freshwater turtle family, Emydidae (pond turtles and terrapins), through histological, pharmacological and immunohistochemical investigations that span several centuries (Bottazzi, 1897; 1906; Fano, 1886; Joyce et al., 2019; 2020; Laurens, 1913; Shaner, 1923; Snyder and Andrus, 1919). Smooth muscle was first identified in *ex vivo* atrial preparations of the European pond turtle (*Emys orbicularis*), distinguishable by its production of characteristic slow tonus-wave contractions that were clearly distinct from the normal rapid contractions of cardiac muscle (Fano, 1886). These tonus contractions were repressed by sympathetic stimulation with adrenaline, and potentiated by vagal stimulation, histamine, and pituitary extract (Bottazzi and Grünbaum, 1899; Dimond, 1959; Gault, 1917; Gruber, 1920; Gruber and Markel, 1918; Sollmann and Rossides, 1927). Early histological studies described a dense layer of smooth muscle lining the luminal side of the atrial wall, originating in the sinus venosus and pulmonary veins, and continuing into the ventricle where its distribution becomes sparse (Laurens, 1913; Shaner, 1923). A recent comparative anatomical study confirmed that atrial smooth muscle is extensive in another emydid turtle, the red-eared slider (*Trachemys scripta*), and is detectable, yet much scarcer, in most other turtles, such as snapping turtles (*Chelydra serpentina*), softshell turtles (*Cyclanorbis senegalensis* and *Pelodiscus sinensis*), and side-necked turtles (*Pelomedusa subrufa*) (Joyce et al., 2020). Atrial smooth muscle is conspicuously absent in terrestrial tortoises, and is not found in other reptiles (Joyce et al., 2020).

Despite these cross-disciplinary approaches, the functional role and evolutionary significance of atrial smooth muscle remains unresolved. In perfused hearts, contraction of the atrial smooth muscle can reduce stroke volume by impeding cardiac filling (Gesell, 1915; Joyce et al., 2019). Hence, atrial smooth muscle has the potential to provide a unique mechanism by which to regulate cardiac output. Because pond turtles exhibit sizeable (two-to three-fold) changes in cardiac output during routine diving behaviour, associated with submergence bradycardia and ventilation tachycardia (Burggren, 1975; Wang and Hicks, 1996), it has been proposed that atrial smooth muscle may play an important role in diving physiology (Joyce et al., 2019; 2020; Joyce and Wang, 2020). During underwater apnoea, when increased vagal tone decreases heart rate (Burggren, 1975), the prolonged cardiac filling time, which would otherwise be expected to increase stroke volume (Joyce et al., 2018), may be compensated by contraction of atrial smooth muscle to reduce cardiac filling and prevent overexpansion (Joyce and Wang, 2020). The precise control of cardiac output during diving facilitates the temporal adjustment of pulmonary ventilation and perfusion matching, increasing or reducing the efficiency of gas exchange based on physiological needs (Malte et al., 2016). While it has been speculated that smooth muscle plays a role in the diving physiology of turtles, for this hypothesis to be supported its presence would be expected to be widespread in diving Testudines, and more extensive in those with enhanced diving capabilities such as sea turtles.

Inclusion of marine turtles, therefore, is essential to examine any putative role in diving, as sea turtles (Chelonioidea) are well adapted for diving and able to remain underwater for several hours and performing dives of greater depth and duration than seen in many other air-breathing vertebrates, including freshwater turtles (Lutcavage and Lutz, 1996). Loggerhead sea turtles, *Caretta caretta*, have been documented on voluntary dives recorded at depths of over 340 m (Narazaki et al., 2015) and for durations of up to 10 h (Broderick et al., 2007; Hawkes et al., 2007; Hochscheid et al., 2007). Interestingly, histamine, a potent constrictor of atrial smooth muscle in pond turtles (Joyce et al., 2019), has been shown to constrict the pulmonary arterial sphincter in loggerhead sea turtles (García-Párraga et al., 2018), which may contribute to the low pulmonary blood flow that limits pulmonary perfusion and gas exchange during diving, *i.e.* the ‘right-to-left shunt’ (Wang and Hicks, 1996). In a recent study, diving bradycardia was also shown to be primarily mediated by the parasympathetic nervous system in loggerhead sea turtles (Saito et al., 2022). It was also previously suggested that this protective vagal diving reflex (i.e. right-to-left cardiac shunt during breath-holding) is disrupted when entangled by-catch turtles become stressed, stimulating and inhibiting the sympathetic and parasympathetic nervous systems respectively, leading to an increased nitrogen uptake and subsequent gas embolism after surfacing (García-Párraga et al., 2014). This protective dive response has also been shown to be interrupted in struggling green sea turtles, *Chelonia mydas*, during involuntary forced dives (Berkson, 1967). Understanding the mechanisms of cardiac regulation, including the potential role for endocardial smooth muscle in sea turtles, is particularly pertinent given that effective regulation of cardiac output during diving is likely to affect the susceptibility to gas embolism (Robinson et al., 2021). The only previous insight into the possible presence of atrial smooth muscle in sea turtles was provided by Bottazzi (1906), who briefly remarked on tonus waves produced by a right atrial *C. caretta* preparation; however, it is not possible to discern if this represented a reproducible observation.

Together, the available literature suggests that, when present, cardiac smooth muscle under parasympathetic control could play a functional role as part of the protective vagal diving reflex in turtles (Joyce et al., 2019; Joyce and Wang, 2020). Based on these presumed roles in diving physiology, we hypothesised that atrial smooth muscle would be extensive in sea turtles, given their extraordinary dive capacity which involves vagal bradycardia (Saito et al., 2022). In order to test this hypothesis and gain insight into the functional significance of atrial smooth muscle, we determined if smooth muscle is present in the loggerhead sea turtle using immunohistochemistry and histology.

## 2. Materials and Methods

### Sample acquisition

Seven loggerhead sea turtle hearts, *Caretta caretta*, (body mass: 6.3 ± 3.5 kg (mean ± SD), range: 1.5 - 12 kg) were obtained from deceased individuals received through the sea turtle stranding network at the Valencian region, operated by the Fundación Oceanogràfic de la Comunitat Valenciana (Valencia, Spain) under the formal approval of the Conselleria de Agricultura, Desarrollo Rural, Emergencia Climática y Transición Ecológica of the Generalitat Valenciana. This included samples from six female and one male turtle; no noticeable sex differences were observed. No animals were euthanized for the purpose of this study; hearts were obtained opportunistically from animals that had recently died or been euthanized due to severe injuries sustained from fisheries interactions along the Valencian coast. Hearts were fixed for 72 - 96 hr in 10% neutral buffered formalin and thereafter stored in 70% ethanol. The fixed hearts were embedded in paraffin wax (Paraplast, Sigma P3558) and sectioned serially into 5 μm transverse or coronal tissue sections using the Leica X Multicut semi-automated rotary microtome (Leica Biosystems, Wetzlar, Germany). Additional samples from juvenile *T. scripta* (n=5, archived at the University of Manchester in paraffin blocks) were included as a positive control group to ensure the protocol was optimised, as a clear atrial smooth muscle signal has been previously characterised in this species (Joyce et al., 2020). These emydid pond turtles had been euthanized in accordance with Schedule 1 of the Animals (Scientific Procedures) Act 1986 as part of an unrelated series of experiments and the hearts had been fixed in 10% neutral buffered formalin prior to embedding in paraffin wax. The embedding orientations of the isolated atria were unknown, however smooth muscle can be clearly identified in both transverse and coronal planes in emydid turtles (Shaner, 1923). Tissue sections were transferred and set onto standard (25 x 75 x 1 mm) or large (52 x 76 x 1 mm) microscope slides.

### Immunohistochemistry

A standard immunohistochemistry protocol was carried out as described previously (Jensen et al., 2016; 2017; Joyce et al., 2019). Detection of smooth muscle was achieved using a mouse monoclonal primary antibody to smooth muscle actin (SMA, Sigma A2547, diluted to 1:600) and a donkey anti-mouse secondary antibody conjugated to the fluorophore Alexa 555 nm (Invitrogen A31570, 1:250). Cardiac muscle was detected with a rabbit recombinant polyclonal primary antibody to cardiac troponin I (cTnI, Invitrogen, 1HCLC, 1:600) and a donkey anti-rabbit antibody conjugated to the fluorophore Alexa 647 nm (Invitrogen A31573, 1:250). Fluorescent imaging was conducted on the 3D-Histech Pannoramic-250 microscope slide-scanner, using the TRITC and CY5 filter sets. Images were taken using the SlideViewer Software (3D-HISTECH).

### Histology

The mounted sections were de-paraffinized in xylene (Sigma Chemical Company, MO, USA), washed twice with ethanol, rehydrated through a graded series of ethanol solutions, placed into water and stained with the following stains: (1) Masson’s trichrome stain, which produce red keratin and muscle fibres, and blue collagen or connective tissue (2) Miller’s elastic stain with Van Geisen’s counterstain (previously employed by Shaner (1923) to detect atrial smooth muscle in emydid turtle hearts), where elastic fibres are in blue/purple, collagen and muscle in red and cytoplasm in yellow. Smooth muscle cells should be distinguishable from cardiac muscle by the endomysium *i.e.* smooth muscle connective tissue. Slides were imaged with bright-field microscopy either using the Olympus BX63 Upright colour camera microscope for large slides or using the 3D-HISTECH Pannoramic-250 microscope slide-scanner for small slides, where snapshots of the slide-scans were taken using the Case Viewer software (3D-HISTECH).

### Statistical Analysis

To analyse the relative proportions of cardiac muscle (red) and smooth muscle (green) in images from the immunohistochemically treated sections, we quantified the relative pixel contributions of each channel in a composite fluorescent image. The analysis was carried out in the ‘QuantCentre’ function of SlideViewer (3D-HISTECH Software). The area to be analysed, *i.e.* the atrium, on each image was defined by creating a closed polygon using the ‘annotation tool’ to outline the atria, and the image segmentation module ‘HistoQuant’ was used to split the red (cTnI) and green (SMA) channels (Fig.6). The area of each colour channel was used to quantify the relative amount of smooth and cardiac muscle as a percentage of total muscle area. Only samples in which the whole atria could be clearly viewed and defined were included in the analysis. Samples in which the tissue had folded or creased, or samples which had high levels of autofluorescence (that interferes with the detection and analysis of the specific smooth muscle signal) were excluded from the analysis. Large slides were unable to be processed for immunohistochemistry. Quantitative analysis was performed following immunohistochemistry on *C. caretta* (n = 5) and *T. scripta* (n = 5). An independent samples t-test was performed in R-Studio software (R Development Core Team, 2013). Data are presented as means ± SD.

## 3. Results & Discussion

A quantitative comparison between *C. caretta* (2.34 ± 1.4% (mean ± SD)) and *T. scripta* (23.3 ± 4.3% (mean ± SD)) revealed a much greater relative area of atrial smooth muscle as a percentage of total muscle area in *T. scripta* (unpaired two-tailed T-test, unequal variance assumed, *p* < 0.0001, t=10.3, df=8) (Fig. 1). Indeed, fluorescent immunohistochemistry and histological staining failed to identify a clear layer of smooth muscle in the atria of loggerhead turtle hearts (Fig. 1), and this was the case for all sections in all hearts (n=7), even those that could not be included in the quantitative analysis. The smooth muscle signals that were detected in *C. caretta* by immunohistochemistry could be largely attributed to trace signals within trabecular bundles, vessel walls of coronary arteries (MacKinnon and Heatwole, 1981), as well as some degree of unavoidable, non-specific binding and co-localisation between muscle types (Supplementary Figure 1). Nevertheless, we were able to verify that our techniques detected clear atrial smooth muscle in the red-eared slider (Fig 1., Supplementary Figs. 2 & 3), where it has previously been characterised in detail (Joyce et al., 2019). Our present quantifications of atrial smooth muscle area in *T. scripta* are similar to the equivalent measurements of an earlier study (17.4 ± 7.9% (mean ± SD)) (Joyce et al. 2020). Vascular smooth muscle was also clearly detected in the coronary arteries and major arteries in the loggerhead turtle (Supplementary Fig. 4), further confirming the validity of the protocols. Whilst in our present comparison, the hearts from *C. caretta* were evidently much larger than those from *T. scripta*, it has previously been shown that the relative amounts of smooth muscle do not change with body mass in *T. scripta* (Joyce et al. 2020) and so differences in the relative amounts of smooth muscle are unlikely to be attributed to differences in body size between these species.

**Figure 1.**
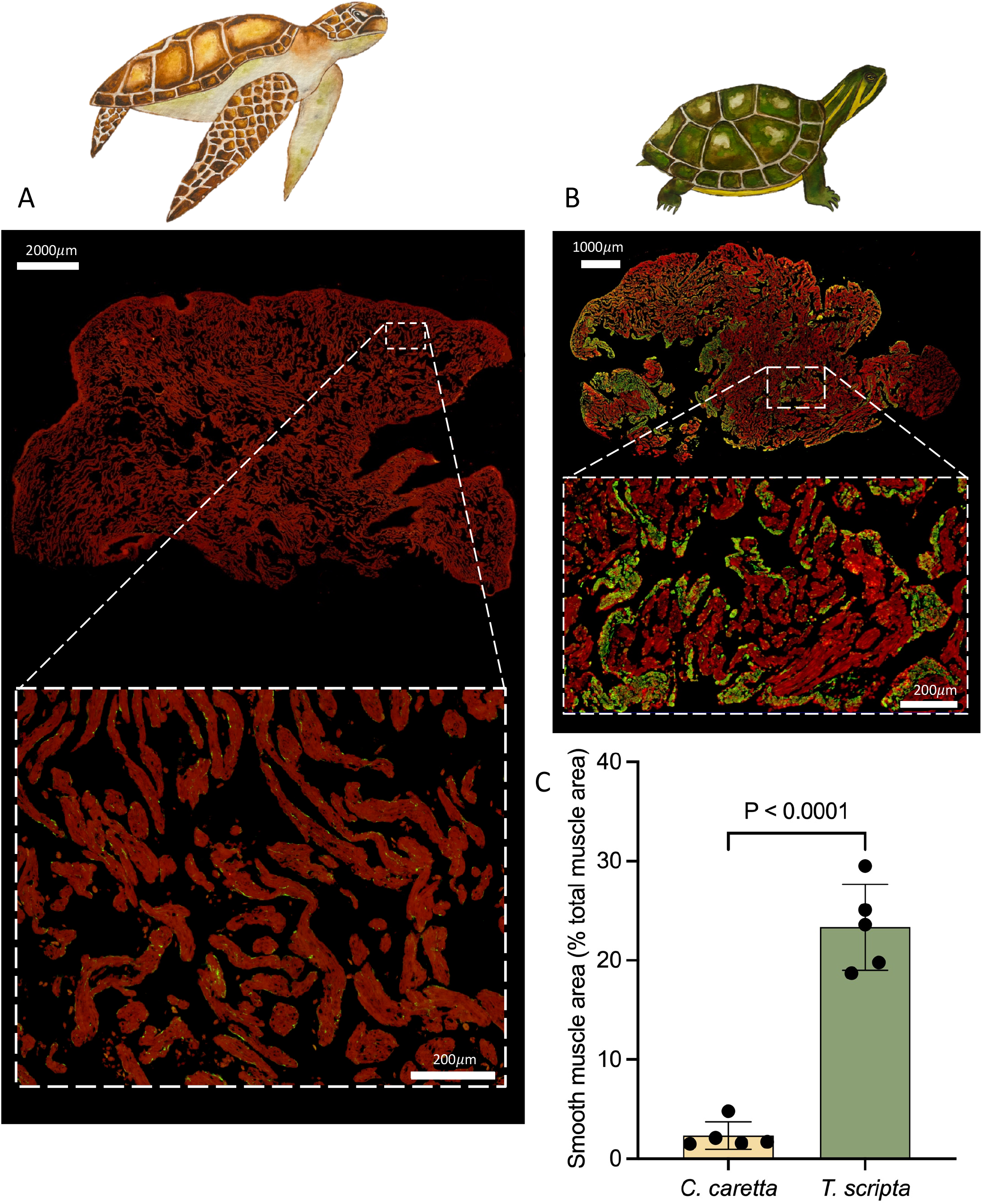
Transverse atrial sections from (A) the loggerhead sea turtle, *Caretta caretta*, and (B) the European pond turtle, *Trachemys scripta* analysed using immunohistochemistry. Smooth muscle and cardiac muscle can be seen respectively in green and red. The presence of smooth muscle on atrial trabeculations was found to be far greater in *Trachemys scripta* (unpaired two-tailed T-test, unequal variance assumed, *p* < 0.0001), and almost completely absent in *C. caretta* (C). Data are means ± SD and individual values.

Although atrial smooth muscle was not detected in loggerhead turtles, smooth muscle was located in the sinus venosus (Figs. 2 & 3, Supplementary Fig. 5), the chamber upstream of the right atrium that connects to the major systemic veins. Transverse sections (Fig. 3; Supplementary Fig. 5) confirm that smooth muscle is found on the luminal side of the sinus venosus_. The presence of smooth muscle in the sinus venosus has been confirmed in eight other species of turtle including side-necked turtles, softshell turtles, snapping turtles, and land tortoises, as well as in amphibians such as the African clawed frog (*Xenopus laevis*) and cane toad (*Rhinella marinus*) (Joyce et al., 2019). The sinus venosus has been shown to function as a contractile cardiac chamber in both *Anolis* lizards and *Python* snakes (Jensen et al., 2017), aiding atrial filling by contracting prior to the atria (Jensen et al., 2014). The proportion of smooth muscle relative to cardiac muscle was found to be significantly greater in the sinus venosus compared to the atria in *T. scripta* (Joyce et al., 2020). We therefore suggest that, in the absence of atrial smooth muscle in *C. caretta*, the sinus venosus, plays an important role in regulating atrial and hence ventricular filling, impeding venous return from the systemic circuit. Together with regulation of pulmonary resistance *via* muscular sphincters in the pulmonary artery (García-Párraga et al., 2018) this may restrict blood-flow to the pulmonary circuit during periods of apnoea whilst diving, facilitating the right-to-left shunt and reducing nitrogen uptake at depth and subsequent risk of decompression sickness after surfacing. The findings presented here, together with the previous work by Joyce et al., (2020), indicate that smooth muscle in the sinus venosus is more highly conserved, and could be functionally more important, than atrial smooth muscle in regulating cardiac output in Testudines. The widespread presence of smooth muscle in the sinus venosus of non-diving Testudines and other reptiles does not detract from a possible role in regulating cardiac output during periods of intermittent lung ventilation, as terrestrial tortoises (*Testudo graeca*) also exhibit intermittent breathing patterns and ventilation tachycardia (Burggren, 1975).

**Figure 2.**
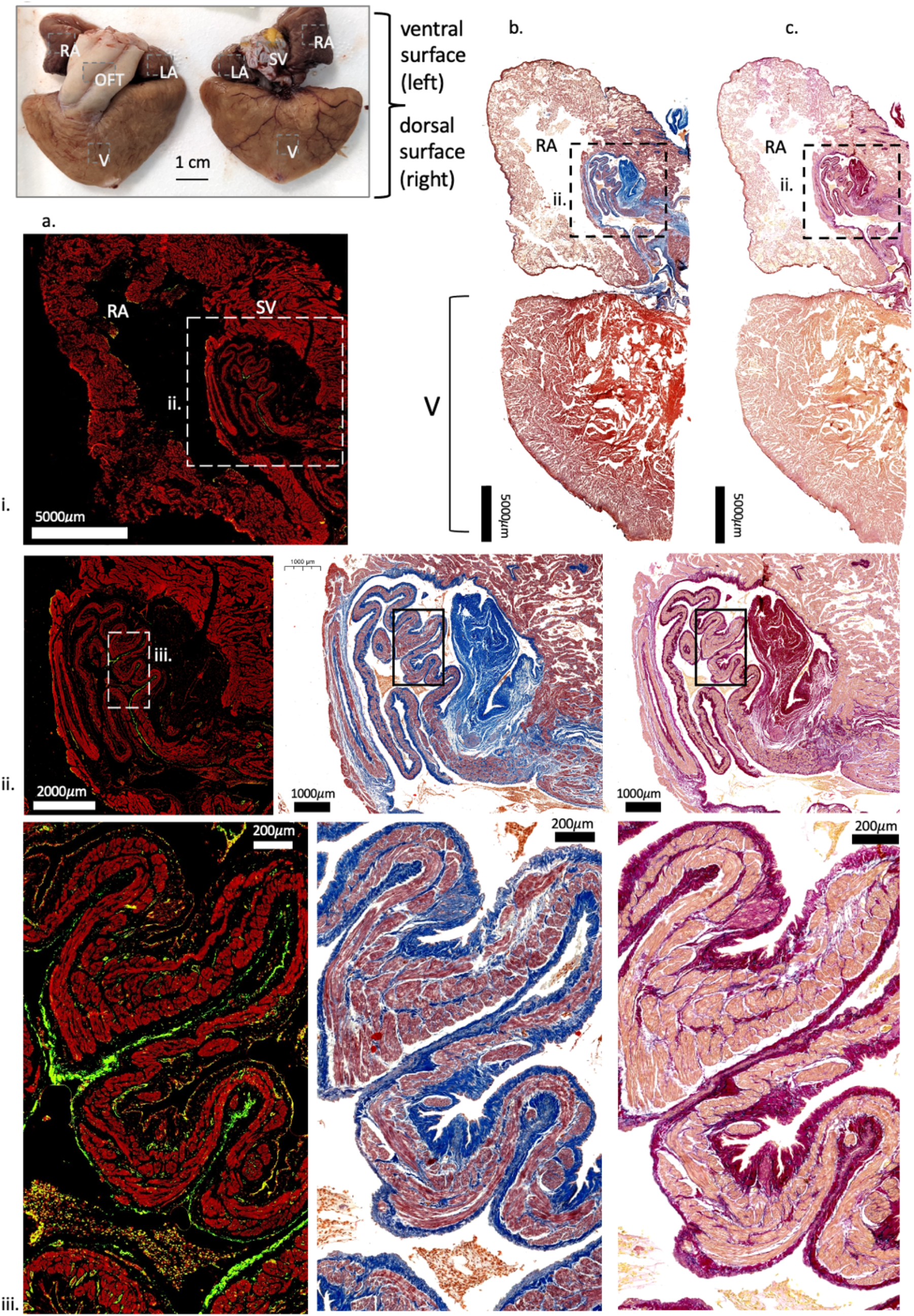
Dorsal coronal cardiac sections of the loggerhead sea turtle, *Caretta caretta*, analysed with immunohistochemistry (column a.) and histology, using a Massons trichrome stain (column b.) and Millers stain with Van Geisons counterstain (column c.). Cardiac chambers are labelled in the top left photograph (for orientation), showing the right atrium (RA), left atrium (LA), sinus venosus (SV), ventricle (V) and outflow tracts (OFT). Smooth muscle and cardiac muscle can be seen respectively in green and red (a.), blue and red/purple (b.) and purple and pink (c.). A distinct layer of smooth muscle was detected in the sinus venosus (row ii.).

**Figure 3.**
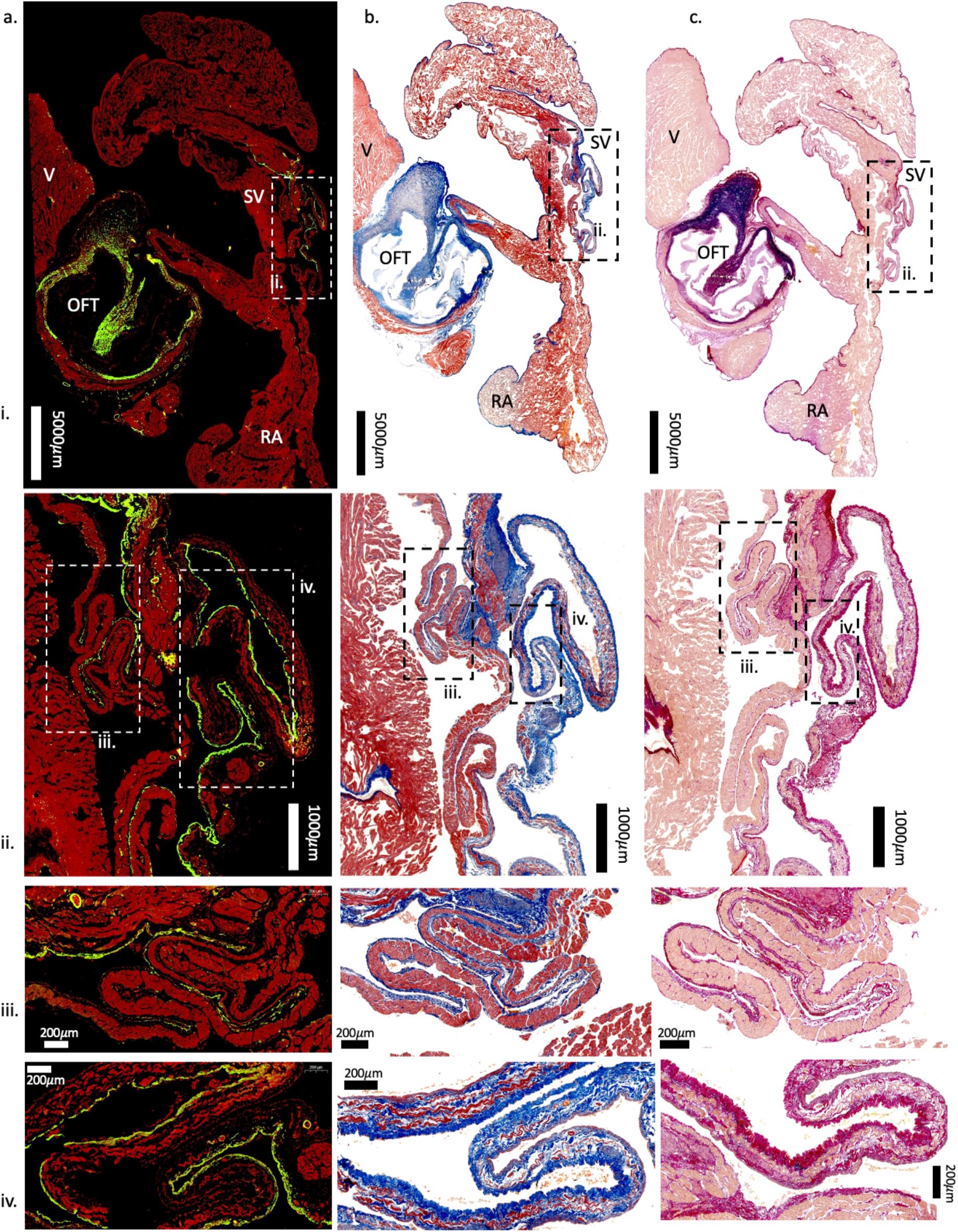
Transverse cardiac sections of the loggerhead sea turtle, *Caretta caretta*, analysed with immunohistochemistry (column a.) and histology, using a Massons trichrome stain (column b.) and Millers stain with Van Geisons counterstain (column c.). Cardiac chambers are labelled showing the right atrium (RA), sinus venosus (SV), ventricle (V) and outflow tracts (OFT). Smooth muscle and cardiac muscle can be seen respectively in green and red (a.), blue and red/purple (b.) and purple/dark pink and pale pink (c.). A distinct layer of smooth muscle was detected in the sinus venosus (row ii.), but not in the atria (i).

The absence of atrial smooth muscle in loggerhead turtles was still nevertheless unexpected, and leads us to reject our hypothesis that it would be extensive in sea turtles that show extraordinary diving capacity. Moreover, our current study undermines the supposition that atrial smooth muscle is an essential component of cardiovascular regulation during diving in turtles generally (Joyce et al., 2019; Joyce and Wang, 2020). The anatomical differences between freshwater pond turtles (extensive atrial smooth muscle) and sea turtles (atrial smooth muscle absent) are interesting given that cardiovascular regulation during diving appears relatively well conserved across the two taxa. For instance, recent research showed that diving bradycardia in loggerhead turtles is mediated by the parasympathetic nervous system (Saito et al., 2022) as in pond turtles (Burggren, 1975). We also expected to find atrial smooth muscle in the loggerhead turtle heart because Bottazzi (1906) earlier referred to tonus waves in right-atrial preparations from the same species. However, a disconnect between anatomy and potentially infrequent tonus waves is not unprecedent; tonus waves were previously reported in the snapping turtle (*Chelydra serpentina*) (Blinks and Koch-Weser, 1963; Pereira, 1924) although these reports were likewise vague and largely anecdotal, and smooth muscle was shown to be very sparse in this species (Joyce et al., 2020). It is also possible that the tonus waves in some of these studies, including the loggerhead turtle, were attributable to the inclusion of some sinus venosus tissue, given that it is in close anatomical association with the right atrium. That atrial smooth muscle is not found in deep diving sea turtles and most other Testudine lineages suggests its reported function in pond turtles need to be further examined. Emydid pond turtles are also renowned for their extreme capacity to tolerate anoxic conditions during overwinter hibernation (Jackson, 2002; Overgaard et al., 2007; Ultsch, 2006), and so it remains possible that atrial smooth muscle plays a functional role in cardiovascular regulation during this extreme condition, although this remains to be fully explored.

Despite their impressive dive capabilities, loggerhead sea turtles have been reported to have fewer adaptations to diving compared to other marine turtles (Williams et al., 2019). During forced submersions, systemic blood flow was found to persist in loggerhead sea turtles (Lutz and Bentley, 1985), whereas in green turtles peripheral vasoconstriction occurred, although this did not persist for the entire submersion (Lutz and Bentley, 1985; Williams et al., 2019). It thus remains possible that loggerhead turtles are not representative of all sea turtles, although it is difficult to envisage the sudden evolutionary loss of atrial smooth muscle expression in one sea turtle species that is maintained in other lineages. Nevertheless, our study unequivocally shows that atrial smooth muscle is not essential in a diving sea turtle species.

In conclusion, atrial smooth muscle is absent in the hearts of *C. caretta*, suggesting that it cannot be involved in the cardiovascular response to diving in sea turtles. Nevertheless, smooth muscle and cardiac muscle were both detected in the sinus venosus, working upstream of the right atrium. In reptiles, contraction of the sinus venosus contributes to atrial filling (Jensen et al., 2017), and the presence of smooth muscle in the sinus venosus is highly conserved across Testudines. We suggest that constriction of sinus venosus smooth muscle impedes venous return and reduces filling of the right atrium, as opposed to being achieved via altering of atrial dimensions through endocardial smooth muscle contraction (Joyce et al., 2019). In the absence of atrial smooth muscle, smooth muscle in the sinus venosus may instead contribute to regulation of cardiac output and pulmonary blood flow in order to minimize nitrogen uptake during diving while still selectively exchanging oxygen and carbon dioxide (García-Parraga et al., 2018, Robinson et al., 2021).

## Data availability

All supporting data is available in the article and its supplementary material.

## Authors’ contributions

WJ conceived the project; all of the authors designed the experiments; LMC carried out the experiments, with the help of WJ and HS. LMC wrote the first draft of the manuscript, with contributions from all authors. All authors gave final approval for publication and agreed to be accountable for all aspects of the content therein.

## Acknowledgments

Thanks to Grace Bako and Joel Rigg from the University of Manchester Histology Core Facility, and to Roger Meadows, Steven Marsden and Peter March in the University of Manchester Bioimaging Facility. Thanks to the Servicio Vida Silvestre of the Conselleria de Agricultura, Desarrollo Rural, Emergencia Climática y Transición Ecológica of the Generalitat Valenciana for the pertinent authorizations for sample collection and to the fishermen collective in the region for all the support bringing by-caught sea turtles to the Oceanogràfic Arca del Mar to attempt rehabilitation. We are also grateful to Laura Cádiz for providing the turtle illustrations used in Figure 1.

