## Supplementary Materials for "Absence of atrial smooth muscle in the heart of the loggerhead sea turtle (*Caretta caretta*): a re-evaluation of its role in diving physiology"

### Supplementary Figures

**Supplementary Figure 1.** Analysis of immunohistochemical atrial samples using ‘QuantCentre’ in ‘SlideViewer’ (3DHISTECH Software) to discriminate between smooth muscle (yellow) and cardiac muscle (blue). (A) *Caretta caretta* cardiac samples included transverse sections (i–iii) and coronal sections (iv) (B) *Trachemys scripta* atrial sample sections (i-v). Blue outlines indicate the regions that were selected for analysis i.e. atria.

**Supplementary Figure 2.** Atrial sections of the Emydid pond turtle, *Trachemys scripta,* analysed under immunohistochemistry (column a.) and histology, using a Masson’s trichrome stain (column b.) and Miller’s stain with Van Geison’s counterstain (column c.). Smooth muscle and cardiac muscle can be seen respectively in green and red (a.), blue/purple and red (b.) and pink/purple and red (c.). A significant amount of smooth muscle was detected in the right atrium of *T. scripta* (first representative individual).

**Supplementary Figure 3.** Atrial sections of the Emydid pond turtle, *Trachemys scripta,* analysed under immunohistochemistry (column a.) and histology, using a Masson’s trichrome stain (column b.) and Miller’s stain with Van Geison’s counterstain (column c.). Smooth muscle and cardiac muscle can be seen respectively in green and red (a.), blue/purple and red (b.) and pink/purple and red (c.). A significant amount of smooth muscle was detected in the right atrium of *T. scripta* (second representative individual).

**Supplementary Figure 4.** Dorsal coronal cardiac sections of the loggerhead sea turtle, *Caretta caretta,* analysed under immunohistochemistry (column a.) and histology, using a Massons trichrome stain (column b.) and Millers stain with Van Geisons counterstain (column c.). Cardiac chambers are labelled showing the right atrium (RA), left atrium (LA), sinus venosus (SV), ventricle (V) and major arteries (MA). Smooth muscle and cardiac muscle can be seen respectively in green and red (a.), blue and red/purple (b.) and purple and pink (c.). A significant amount of smooth muscle was detected in the outflow tracts (row ii.), and only very small amounts of smooth muscle were detected in the atrial trabeculae within the atrial walls (seen in row iv).

**Supplementary Figure 5.** Transverse cardiac sections of two loggerhead sea turtle hearts, *Caretta caretta,* analysed under immunohistochemistry (row a., d.) and histology, using a Massons trichrome stain (row b., e.) and Millers stain with Van Geisons counterstain (row c, f.). Cardiac chambers are labelled showing the right atrium (RA), the left atrium (LA) and the sinus venosus (SV). Smooth muscle and cardiac muscle can be seen respectively in green and red (a., d.), blue and red/purple (b., e.) and dark pink and pale pink (c., f.). A distinct layer of smooth muscle was detected in the sinus venosus, but not in the atrial walls.

Supplementary Figure 1.

A)

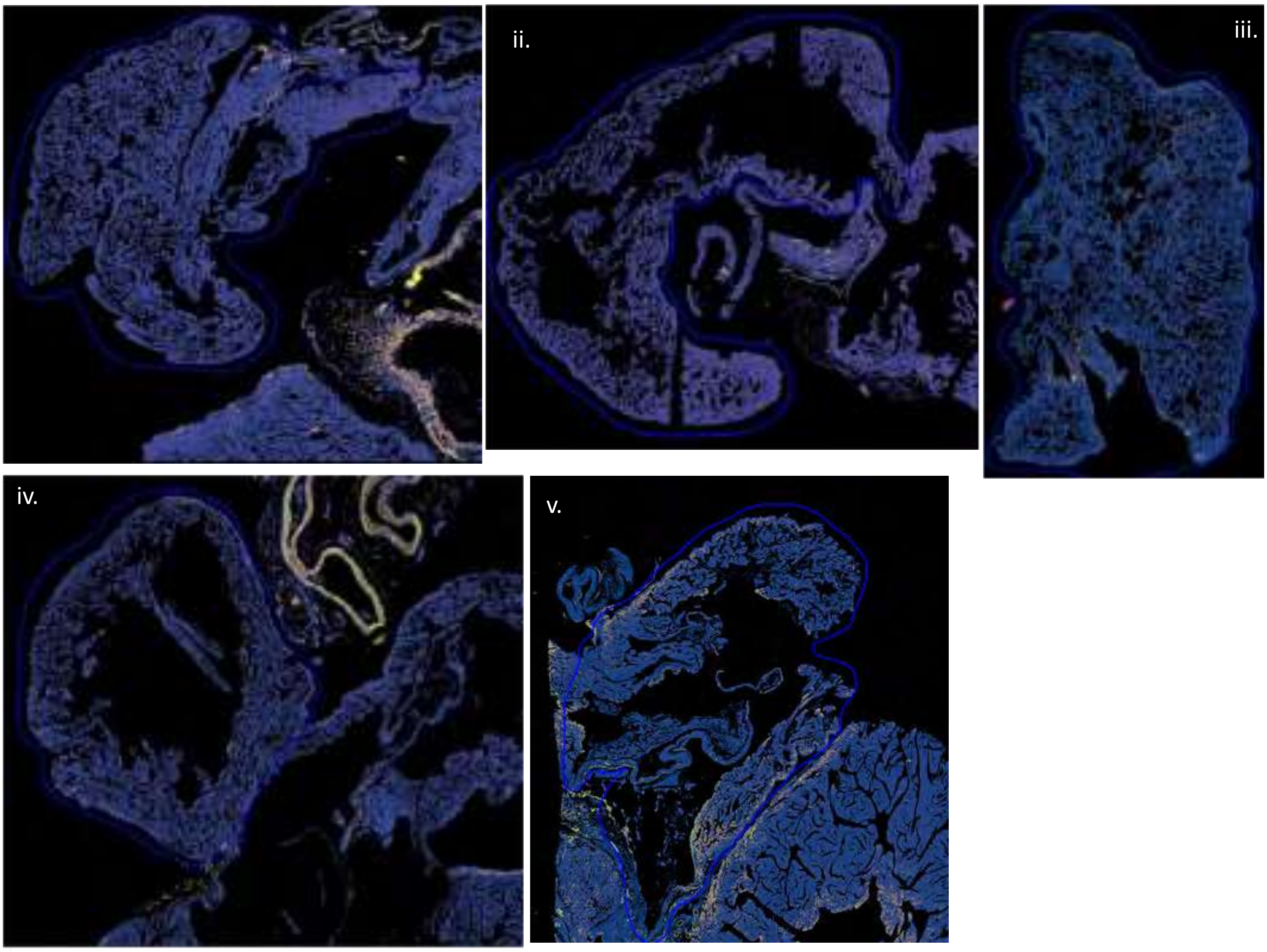


B)

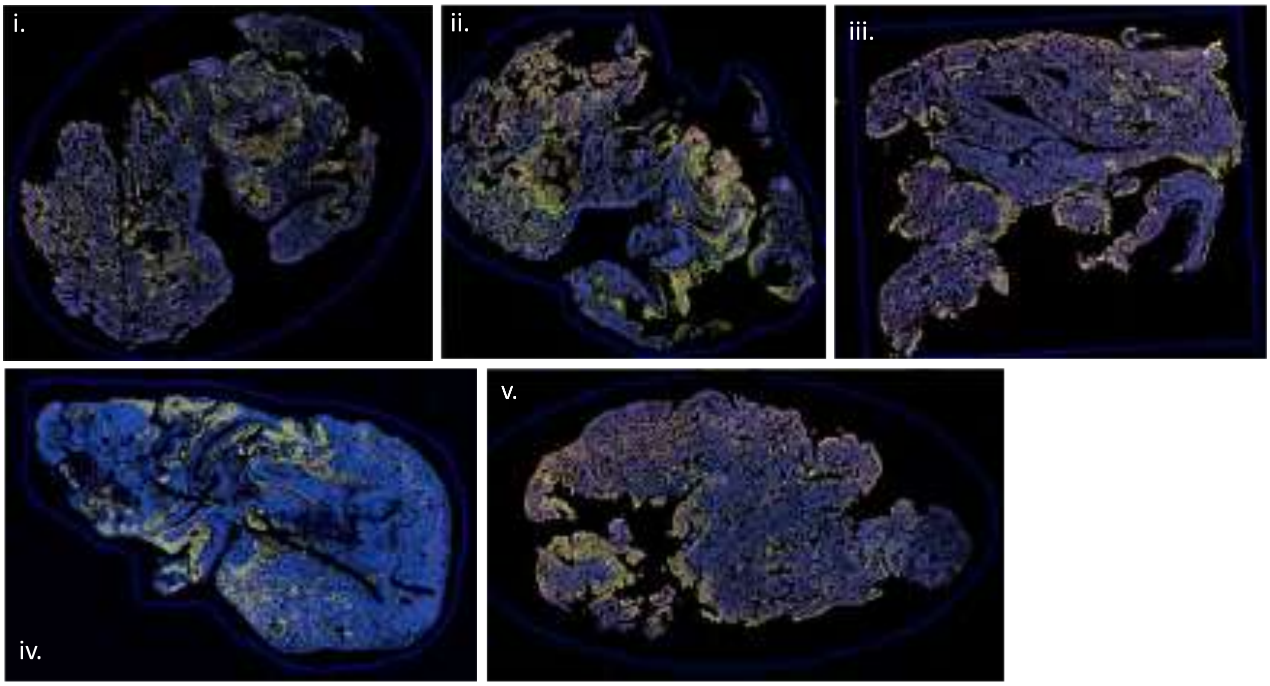

Supplementary Figure 2.

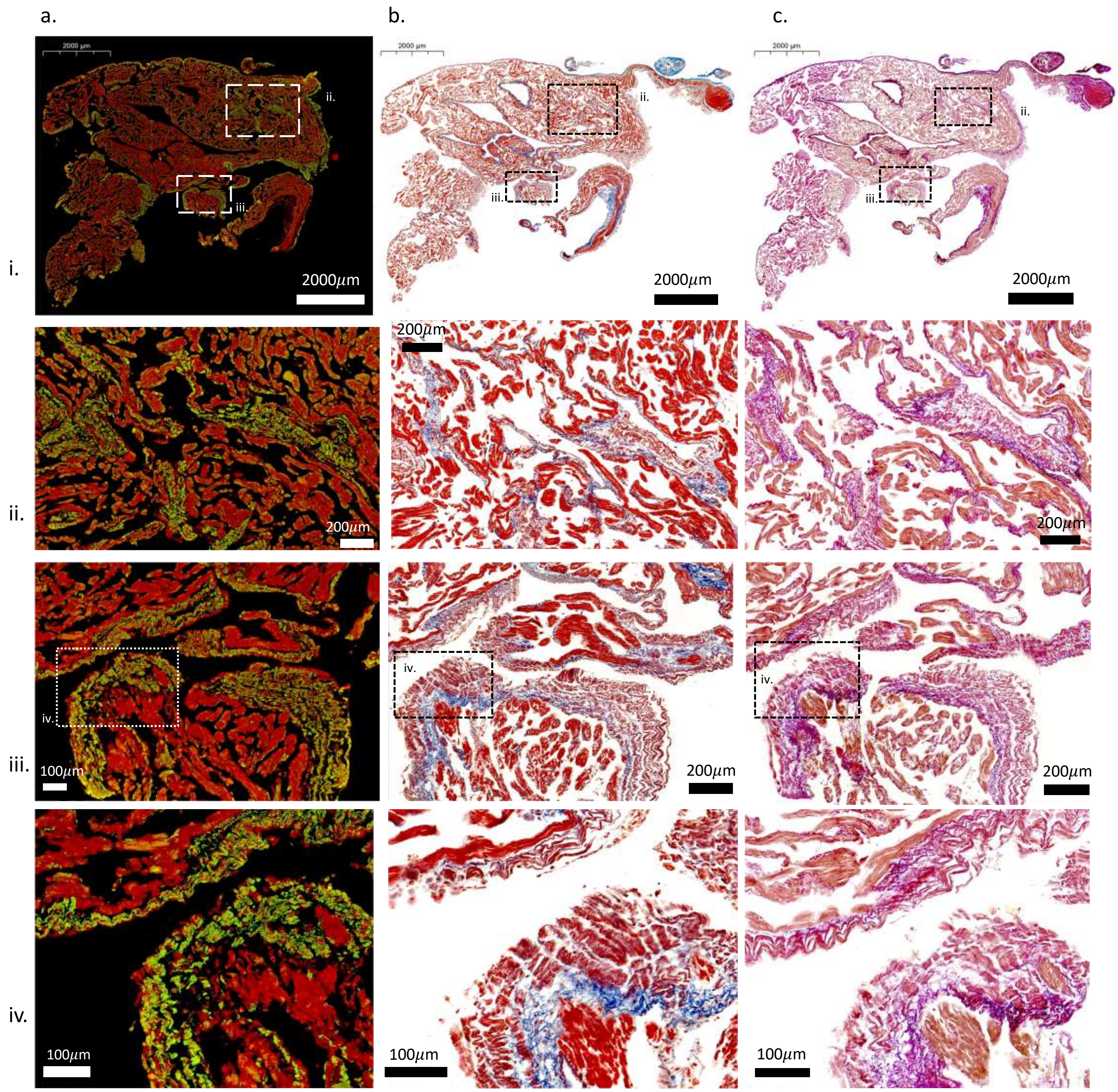

Supplementary Figure 3.

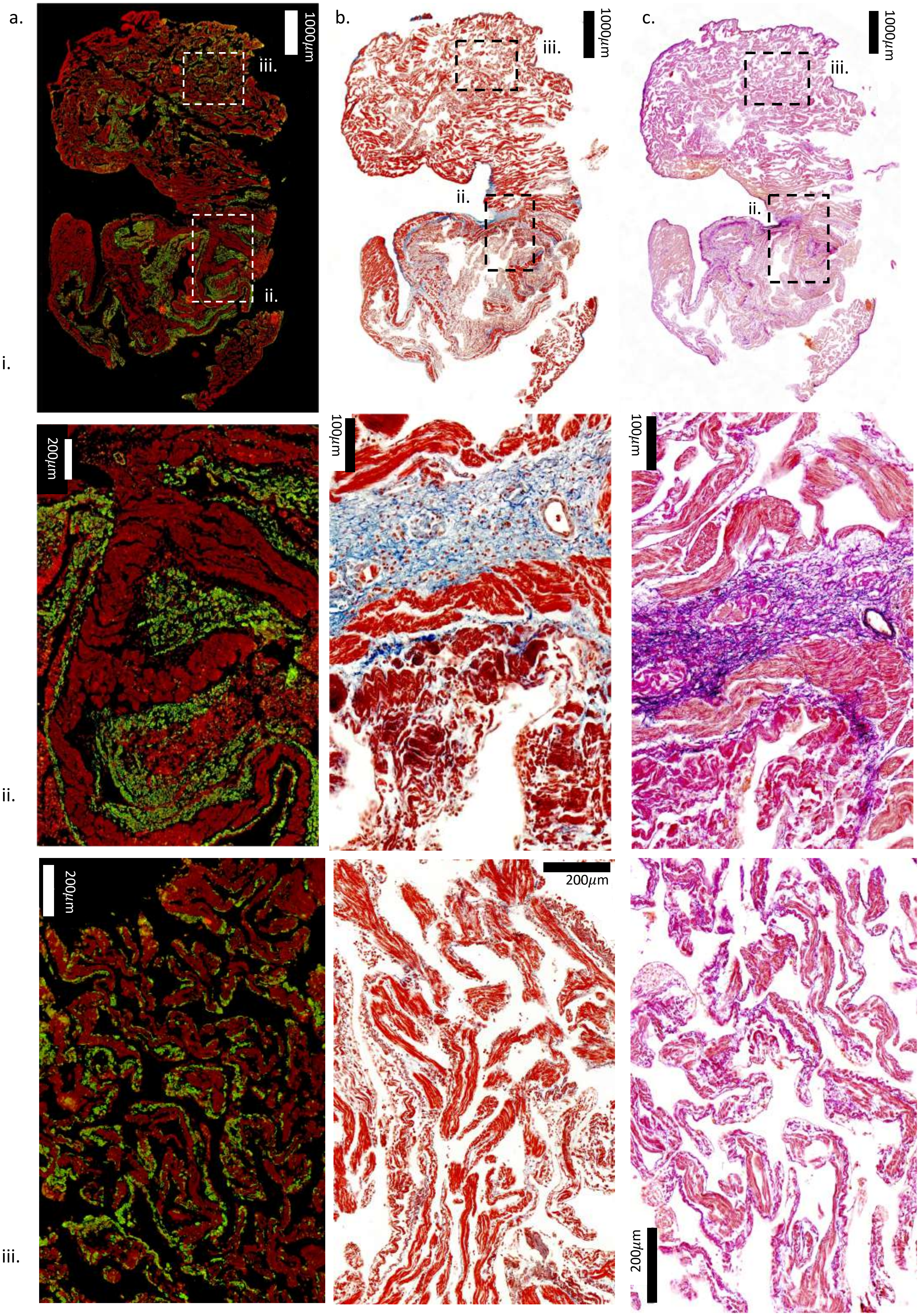

Supplementary Figure 4.

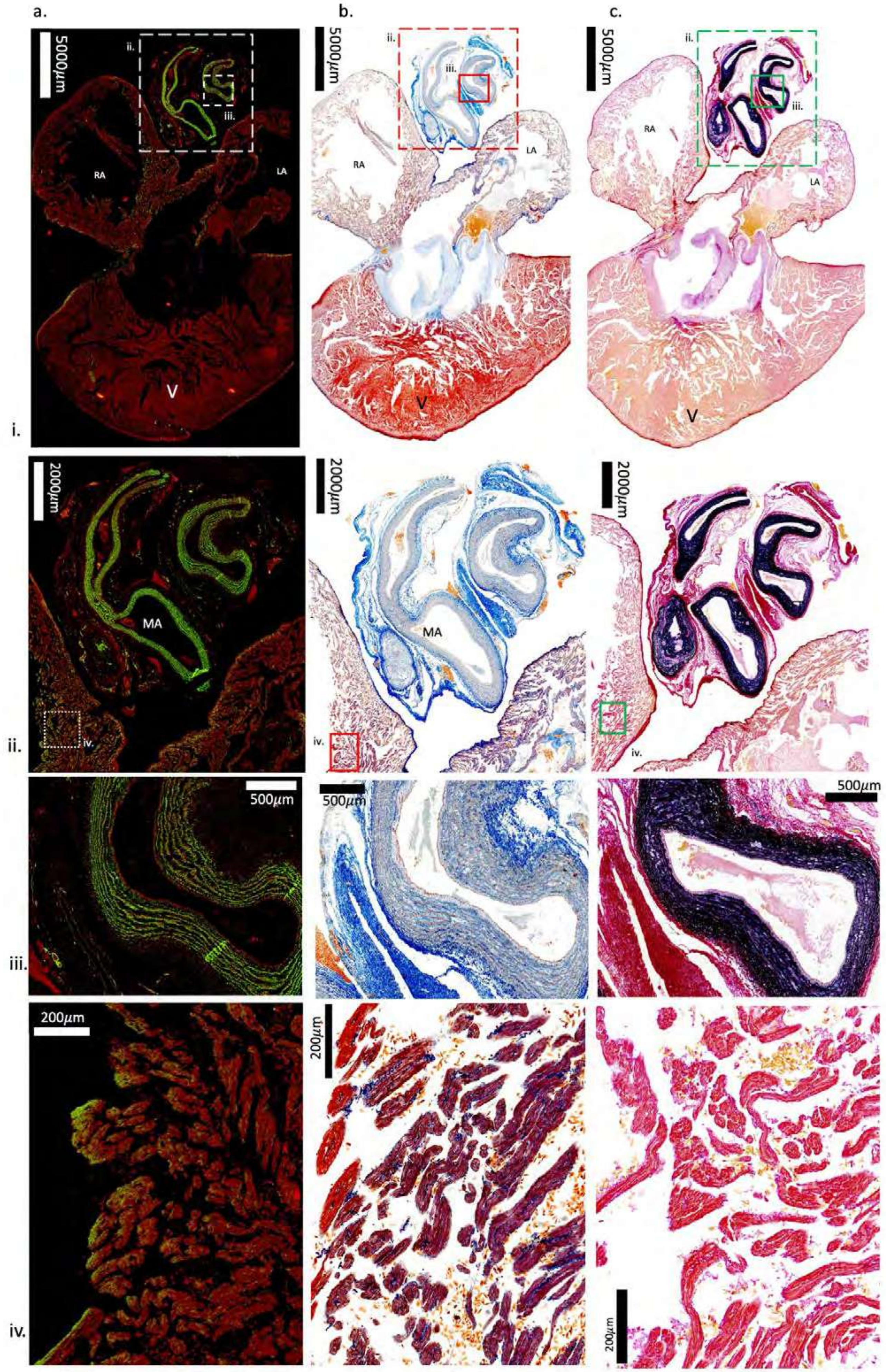

Supplementary Figure 5.

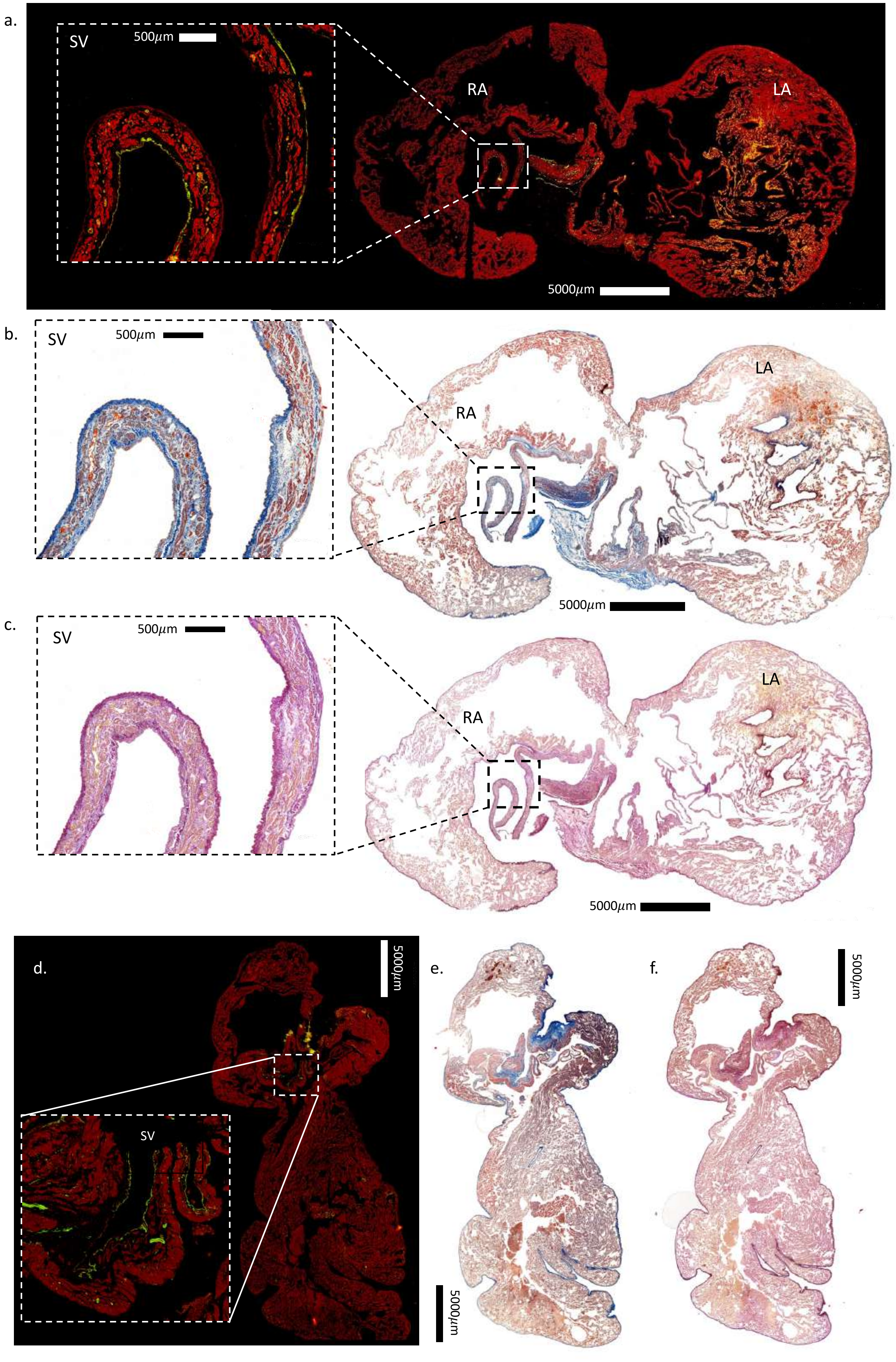

### Supporting Data

**Supplementary Table 1.** Individual values for smooth muscle area (% total muscle area):

| *C. caretta* | *T. scripta* |
|---|---|
| 1.7 | 19.8 |
| 2.1 | 18.7 |
| 4.8 | 23.6 |
| 1.6 | 29.5 |
| 1.5 | 25.1 |
